# CAR-SPLASH identifies nascent pre-mRNA structures implicated in kinetic coupling and alternative splicing

**DOI:** 10.1101/2025.03.02.641068

**Authors:** Hossein Shenasa, Nova Fong, Benjamin Erickson, Ira Iosub, Jernej Ule, David L. Bentley

## Abstract

Pre-mRNA splicing is kinetically coupled to transcription as shown by the widespread effects of transcription speed on alternative splicing outcomes. The molecular basis for such kinetic coupling is incompletely understood, but one potential mechanism is through elongation rate dependent alternative folding pathways of the nascent pre-mRNA. To search for RNA structures in nascent pre-mRNA, we modified SPLASH (1) (<u>S</u>equencing of <u>P</u>soralen Crosslinked, <u>L</u>igated <u>A</u>nd <u>S</u>elected <u>H</u>ybrids) for use with <u>C</u>hromatin <u>A</u>ssociated <u>R</u>NA. We applied this new method called CAR-SPLASH to cells expressing WT and slow mutant RNA polymerase II and identified > 3000 intramolecular RNA structures of which > 400 are proximal to splice sites. Antisense oligonucleotide (ASO) disruption of several such structures that sequester splice sites has a major impact on alternative splicing outcomes, even though the ASOs do not directly disrupt splice sites. ASO disruption of novel regulatory elements that we designate “RNA kinetic switches”, modified alternative splicing of NISCH Exon 18, *GAK* Exon 7 and MEGF8 Exon 14 in a way that depends on the rate of transcription elongation. We propose that these switches mediate kinetic coupling via effects of transcription speed on folding of nascent RNA structures that modulate alternative splicing, and that many nascent RNA structures can thereby serve as new targets for splice modifying ASOs.

**Significance Statement:** Alternative splicing (AS) of mRNAs is a major regulator of gene expression that is frequently disrupted is disease. AS is affected by the speed of RNA polymerase II that synthesizes mRNA precursors, but how such kinetic coupling works is not well understood. We developed a method, CAR-SPLASH, to capture RNA structures in nascent RNA chains by cross-linking with psoralen. CAR-SPLASH identified novel structural regulatory elements we call “RNA kinetic switches” that control AS in a way that depends on the speed of transcription. RNA kinetic switches are new targets for splice modifying antisense oligonucleotides that could have therapeutic value.

## Introduction

Nascent pre-mRNA rapidly folds on itself, which affects its interactions with RNA binding proteins and other regulatory factors, including the spliceosomal snRNPs (2–6). Moreover, transcription is coupled with pre-mRNA splicing both physically and kinetically. Physical coupling is mediated by dynamic interactions between RNA polymerase II (pol II) and splicing factors including SR proteins, U2AF (7–10) and most notably U1 snRNP which directly contacts the Rpb2 and Rpb12 subunits of the polymerase (11). Kinetic coupling between transcription and pre-mRNA splicing is evident from the widespread effects of altering transcription speed on alternative splicing outcomes (12–16) (17). For example, a mutant of the pol II large subunit (Rpb1, R749H) that elongates 3-4X more slowly on average than wild type (WT) pol II affects alternative splicing of thousands of alternative exons and retained introns (17), an effect that can be reproduced with transfected minigenes (16). Notably transcription rate can also affect the outcome of splicing that is completed post-transcriptionally because it is regulated by RNA folding and association with RNA binding proteins (RBPs) that occur co-transcriptionally (18, 19). Physiological variables such as transcriptional activation, DNA damage, GC content of the gene, chromatin modifications, and the regulated activities of pausing and elongation factors can influence pol II speed (20–26) and could thereby modulate splicing outcomes. However, these factors are difficult to control experimentally and thus pol II rate mutants are perhaps the most powerful tool to study kinetic coupling *in vivo* (16). These mutants demonstrate the requirement for an optimal rate of pol II elongation for proper splicing of many alternative exons (17), but the underlying mechanisms that mediate this kinetic coupling are still poorly understood. The classic “window of opportunity” and “first come first served” models postulate that slower elongation promotes alternative exon inclusion by allowing more time for upstream splice sites to be recognized before downstream competing exons are synthesized (16, 27). While these models hold true for certain kinetically regulated alternative splicing (AS) events, they do not account for all effects of transcription speed on AS.

Another potential mechanism for how transcription speed can modulate splicing is through elongation rate-sensitive folding of the nascent pre-mRNA. The rate of transcript elongation can affect co-transcriptional RNA folding pathways by introducing different kinetic traps that change the ensemble of structures formed (19, 28–34). In turn, pre-mRNA secondary structure can influence alternative splicing outcomes by bringing splice sites together sequestering or presenting splice sites, or by changing the accessibility of splicing enhancer and silencer elements that are recognized by RBPs (3, 35–45). However, the documented examples of naturally occurring nascent pre-mRNA structures that are directly responsible for modulating splicing of cellular genes are still quite limited (35, 46–50). Previous studies have predicted functionally relevant pre-mRNA structures based on sequence conservation (51–54). Several global studies of cellular nascent RNA structure have been conducted, but they have uncovered only a limited number of specific structures that control of alternative splicing (19, 55–58). In this report we set out to search for RNA structures that mediate kinetic coupling or influence alternative splicing by taking advantage of cells expressing WT and slow mutant pol II, and by establishing <u>C</u>hromatin <u>A</u>ssociated <u>R</u>NA-<u>S</u>equencing of <u>P</u>soralen crosslinked, <u>Li</u>gated <u>A</u>nd <u>S</u>elected <u>H</u>ybrids (CAR-SPLASH), a global method to map nascent RNA duplexes which builds upon the SPLASH protocol for mRNAs (1). We applied *Tosca* (59) to detect RNA structures implicated in kinetic coupling of transcription elongation to alternative splicing. CAR-SPLASH identified several thousand psoralen crosslinked structures within pre-mRNAs, including several hundred in the vicinity of splice sites. We assessed the function of several such regulatory elements in alternative splicing by targeted ASO disruption. This approach uncovered “RNA kinetic switch” elements in the *NISCH, GAK* and *MEGF8* genes that modulate alternative splicing in a transcription rate-sensitive way through splice site juxtaposition or sequestration. In summary, CAR-SPLASH provides a potent way of investigating the pre-mRNA secondary structure landscape that influences kinetic coupling and alternative splicing.

## Results

### CAR-SPLASH for identification of nascent RNA structures

To investigate the role of secondary structure in kinetic coupling and alternative splicing, we set out to establish a method that identifies specific structures in nascent pre-mRNA by adapting SPLASH (Sequencing of Psoralen Crosslinked, Ligated And Selected Hybrids) method (1), a powerful method to identify RNA duplexes formed *in vivo*. This approach involves purification of psoralen crosslinked RNA followed by proximity ligation and next generation sequencing, to identify the two arms of each RNA duplex and obtain sequence specific base pairing information for local and long-range structures. We adapted SPLASH for use with nascent transcripts and incorporated an improved biotin conjugated psoralen analog, AP3B (60). To enrich for nascent pre-mRNA, we isolated <u>C</u>hromatin-<u>A</u>ssociated <u>R</u>NAs from urea washed nuclei (61) and combined this with the adapted SPLASH protocol in a procedure designated CAR-SPLASH (Figure 1A). To identify nascent RNA structures that are potentially sensitive to the rate of transcript elongation, we applied CAR-SPLASH to HEK293 cells that express α-amanitin resistant WT or slow mutant (R749H) pol II large subunits (Rpb1). We used *Tosca* (59), a Nextflow RNA proximity ligation data analysis pipeline to detect hybrid reads in which two arms separated by a gap correspond to the interacting RNA elements ligated together. Unique molecular identifiers (UMIs) were incorporated in the libraries to allow removal of PCR duplicates (62). Approximately 4.5 and 4.3 million mapped hybrid reads (from two pooled replicates Supplementary Figure S1) were obtained for the WT and slow pol II mutant cells, respectively. To delineate unique duplexes, *Tosca* performs a stringent graph-based clustering of the hybrid read alignments, retaining only those supported by at least two hybrid reads overlapping by at least 50% in both arms (see Methods). Approximately 60% of all intramolecular structures were intronic, indicating nascent RNA enrichment (Figure 1B). We focused on intramolecular structures in pre-mRNAs and lncRNAs with RNAse III digested gaps longer than 3 bases, and identified ∼2,900–3,600 unique structures in WT and the slow pol II mutant of which 484 were shared (Figure 1C, Table S1). Many structures (843 WT and 587 R749H, 178 shared, Table S1) identified by CAR-SPLASH are proximal to those predicted on the basis of sequence conservation (53)(Figure 1C, Table S1). We also detected many cross-linked intermolecular structures which were not analyzed further here (Supplementary Figure 1C).

**Figure 1.**
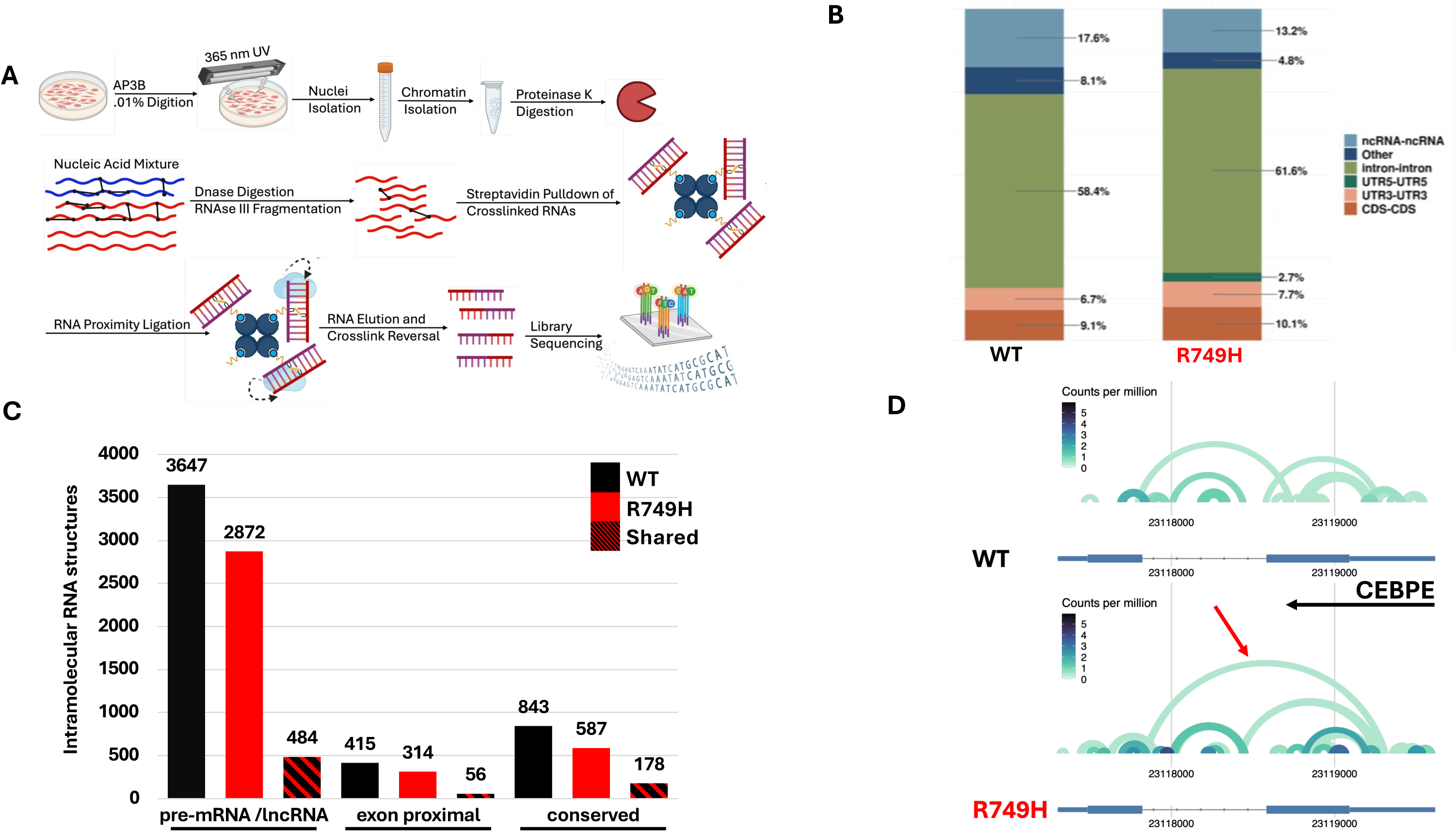
CAR-SPLASH identifies psoralen cross-linked RNA structures in nascent pol II transcripts. **A.** Overview of CAR-SPLASH method. **B.** The distribution of intramolecular RNA structures in different genomic regions in the WT or slow (Rpb1 R749H) pol II mutant. **C.** Intramolecular pre-mRNA and lncRNA structures detected by CAR-SPLASH in WT and slow pol II conditions (columns 1-3). Structures within 200 nucleotides of an exon (columns 4-6) and those proximal to conserved RNA structures (54)(columns 7-9) are shown. **D.** Arc plots of RNA structures detected by CAR-SPLASH in the CEBPE primary transcript. Red arrow indicates a structure exclusively detected in the slow mutant.

Because of the limited sensitivity of RNA proximity ligation methods, combined with the high stringency of the clustering algorithm (59) we used to delineate duplexes, the catalogue of nascent RNA structures we identified is far from complete and we are not able to determine if coverage differences between WT and slow pol II samples are significant. Targeted approaches (35, 63–65) are usually required to obtain the read depth required to identify alternative candidate structures within an ensemble. However, in a few cases such as the CEBPE pre-mRNA CAR-SPLASH yielded sufficient coverage to identify multiple mutually exclusive RNA structures that might have regulatory significance (Figure 1D).

### A pre-mRNA structure that controls alternative exon inclusion

To identify RNA structures that could potentially impact alternative splicing we selected those where at least one arm is in an intron within 200 bases of an exon. This analysis identified 415 and 314 exon proximal pre-mRNA structures in WT and slow pol II mutant cells respectively, of which 56 are shared (Figure 1C, columns 4-6). These results suggest that many of the intronic RNA structures captured by CAR-SPLASH could potentially influence alternative splicing and merit further study.

To investigate whether structures that form in the vicinity of splice sites actually affect splicing we disrupted them with antisense 2’-O-methoxyethyl modified RNA oligonucleotides (ASOs). Initially, we interrogated a structure detected in WT cells that is predicted to sequester the 5’ splice site (ss) of BRSK2 exon 19 (Figure 2A, Supplemental Figure1A). A specific ASO targeting the downstream arm of the RNA duplex distal from the splice site or an irrelevant control oligonucleotide (see Methods) was transfected into WT and slow pol II mutant HEK293 cells in triplicate and splicing was monitored by RT-PCR. The BRSK2 specific ASO results in a small but reproducible increase in inclusion of exon 19 in both WT and slow pol II mutant cells (Figure 2B,C). We conclude that the pre-mRNA structure detected by CAR-SPLASH functions to sequester the 5’ ss of *BRSK2* exon 19 and enhance skipping of this exon. ASO disruption of the structure presumably makes this 5’ ss more accessible resulting in increased exon inclusion.

**Figure 2.**
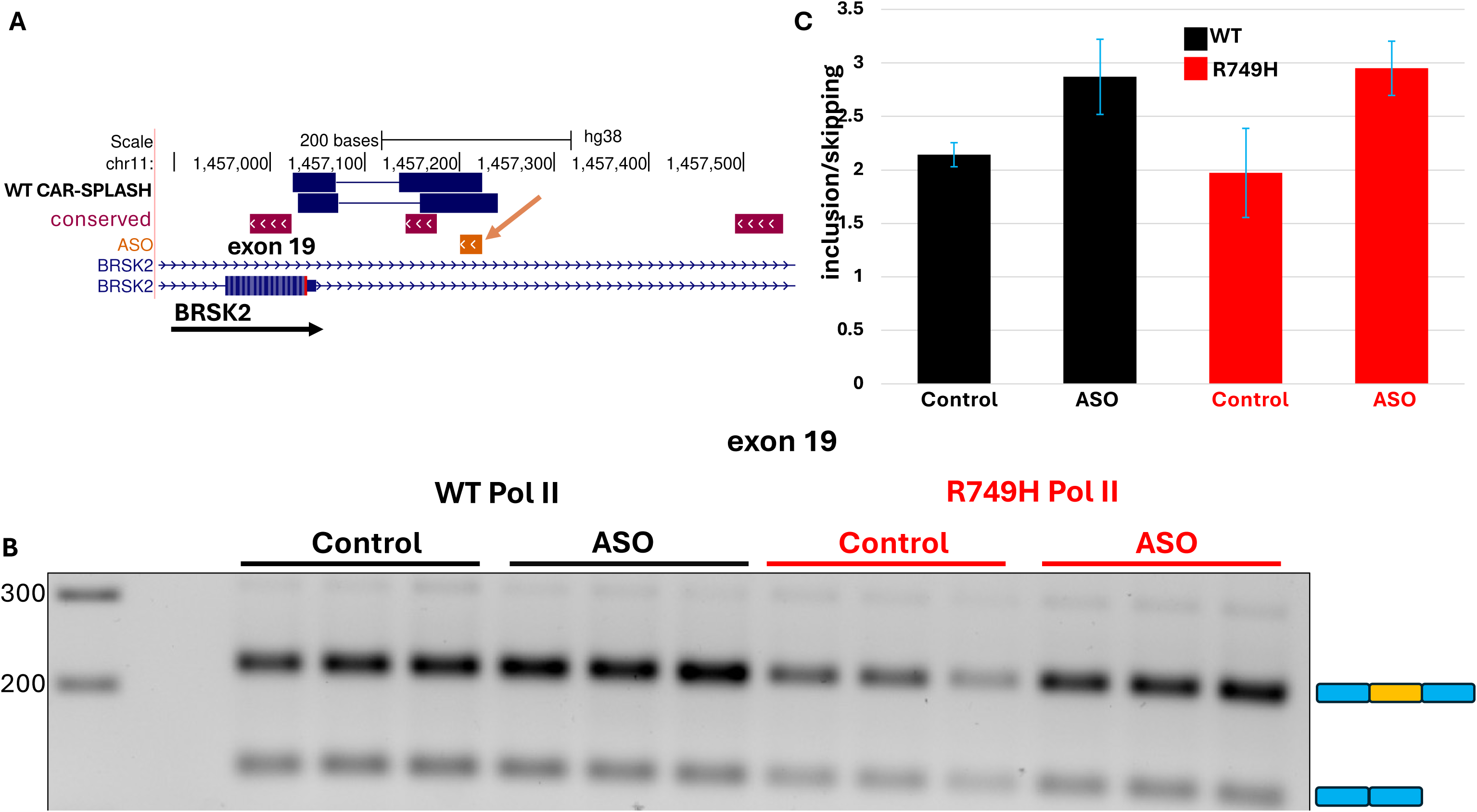
CAR-SPLASH identifies a structure that regulates BRSK2 exon 19 skipping through 5’ ss sequestration. **A.** UCSC genome browser view of CAR-SPLASH reads detected in WT pol II expressing cells (black). The ASO targeting the downstream arm of the RNA structure (see Supp. Fig. 2A) is indicated (orange arrow). **B.** RT-PCR analysis of BRSK2 exon 19 inclusion/skipping in cells expressing WT or R749H slow pol II transfected with the BRSK2 specific ASO or a control oligonucleotide (3 biological replicates). **C.** The average ratio of exon 19 inclusion to skipping. Note enhanced exon inclusion when the structure predicted to sequester the 5’ss is disrupted by the specific ASO.

### RNA kinetic switches: Rate sensitive structures that influence alternative splicing

To discover RNA structures that might play a role in kinetic coupling of transcription with splicing, we intersected (see Methods) the positions of RNA structures with rate-sensitive splicing events detected by rMATS (66). For this analysis, we repeated the RNA-seq of HEK293 cells expressing WT and slow pol II (Rpb1 R749H) (17) at greater read depth using rRNA depletion instead of polyA+ selection, and identified ∼20,000 significantly affected (FDR < 0.05) alternative splicing events (skipped exons, retained introns, alternative 5’ splice sites, alternative 3’ splice sites and mutually exclusive exons) including ∼8300 increased exon skipping and ∼ 4700 increased exon inclusion events in the slow mutant (Figure 3A). We uncovered ∼200 RNA structures in WT and slow pol II expressing cells that form within 200 bases of a rate sensitive alternative splicing event (Figure 3B) and therefore might be involved in regulation of this process. Several structures were selected for further study on the basis that they are predicted to either juxtapose or sequester splice sites. We first investigated a structure detected by CAR-SPLASH in both the WT and slow pol II mutant that partially sequesters the 5’ ss of *NISCH* exon 18 and also bridges across most of the downstream intron potentially juxtaposing the 5’ and 3’ ss (Figure 4A). We targeted the downstream arm of the RNA duplex located 28-48 bases from the 3’ ss with an ASO (Supplemental Figure 2B). Remarkably, disruption of this structure caused skipping of exon 18 and this effect is stronger in the slow pol II mutant than in WT (Figure 4B, C). These results suggest that the intronic RNA structure downstream of exon 18 functions to enhance splicing of intron 18 presumably by promoting cross-intron contacts with consequent inclusion of exon 18. Furthermore, this RNA structural enhancer of splicing appears to have a greater effect under conditions of slow pol II elongation, possibly because its formation is favored under those conditions.

**Figure 3.**
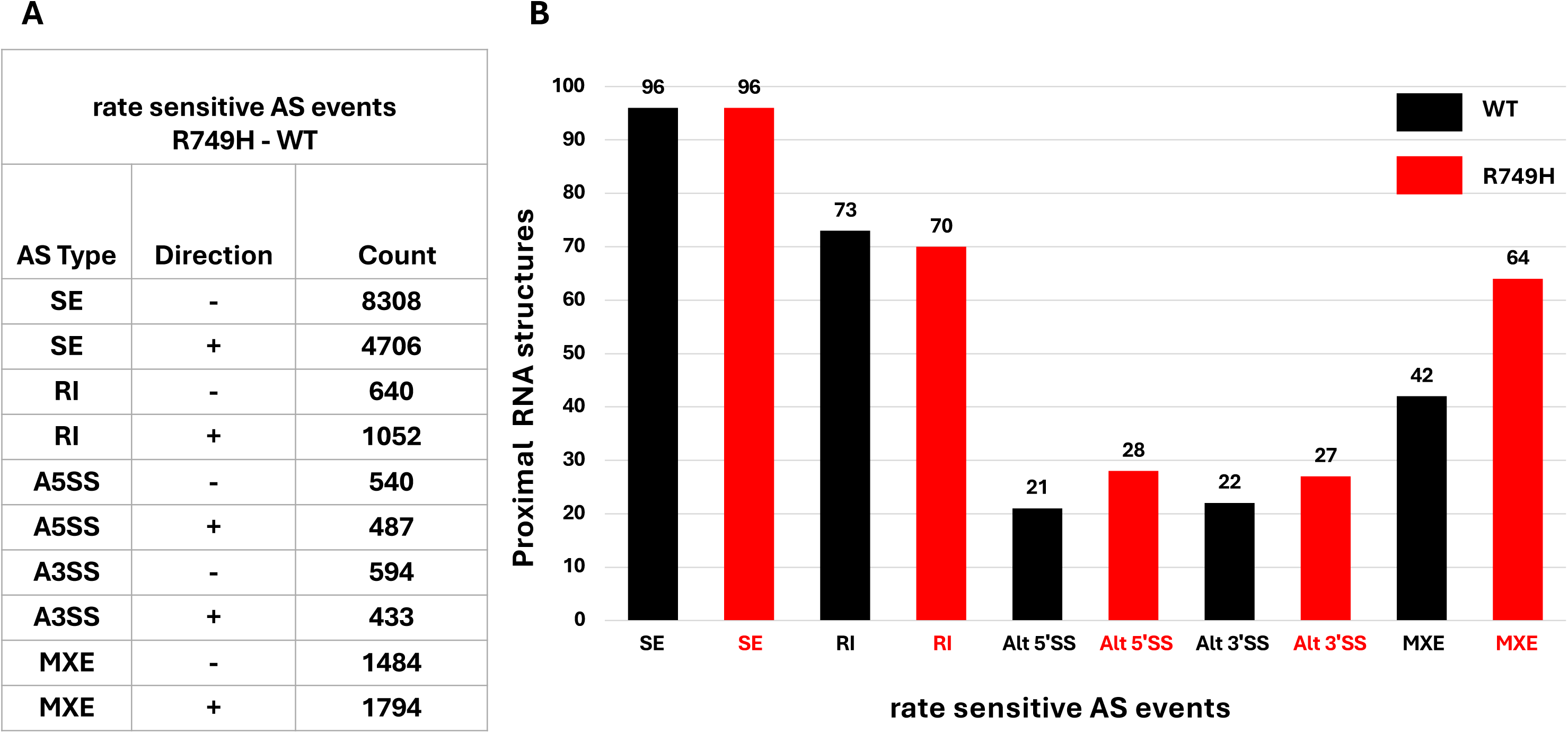
RNA structures proximal to elongation rate sensitive splicing events. **A.** Transcription elongation rate sensitive alternative splicing events detected by rMATS analysis (3 replicates p-value < 0.05 and FDR < 0.05) of RNA-seq data from cells expressing WT and slow mutant pol II (R749H). (SE skipped exon, RI retained intron, MXE mutually exclusive exon, A3SS and A5SS, alternative 3’ and 5’ splice site, Direction: - and + refers to differences between R749H-WT. i.e. SE: – indicates more inclusion in WT than R749H and + indicates more inclusion in R749H than WT. **B.** CAR-SPLASH identified RNA structures located within 200 bases of rate sensitive alternative splicing events.

**Figure 4.**
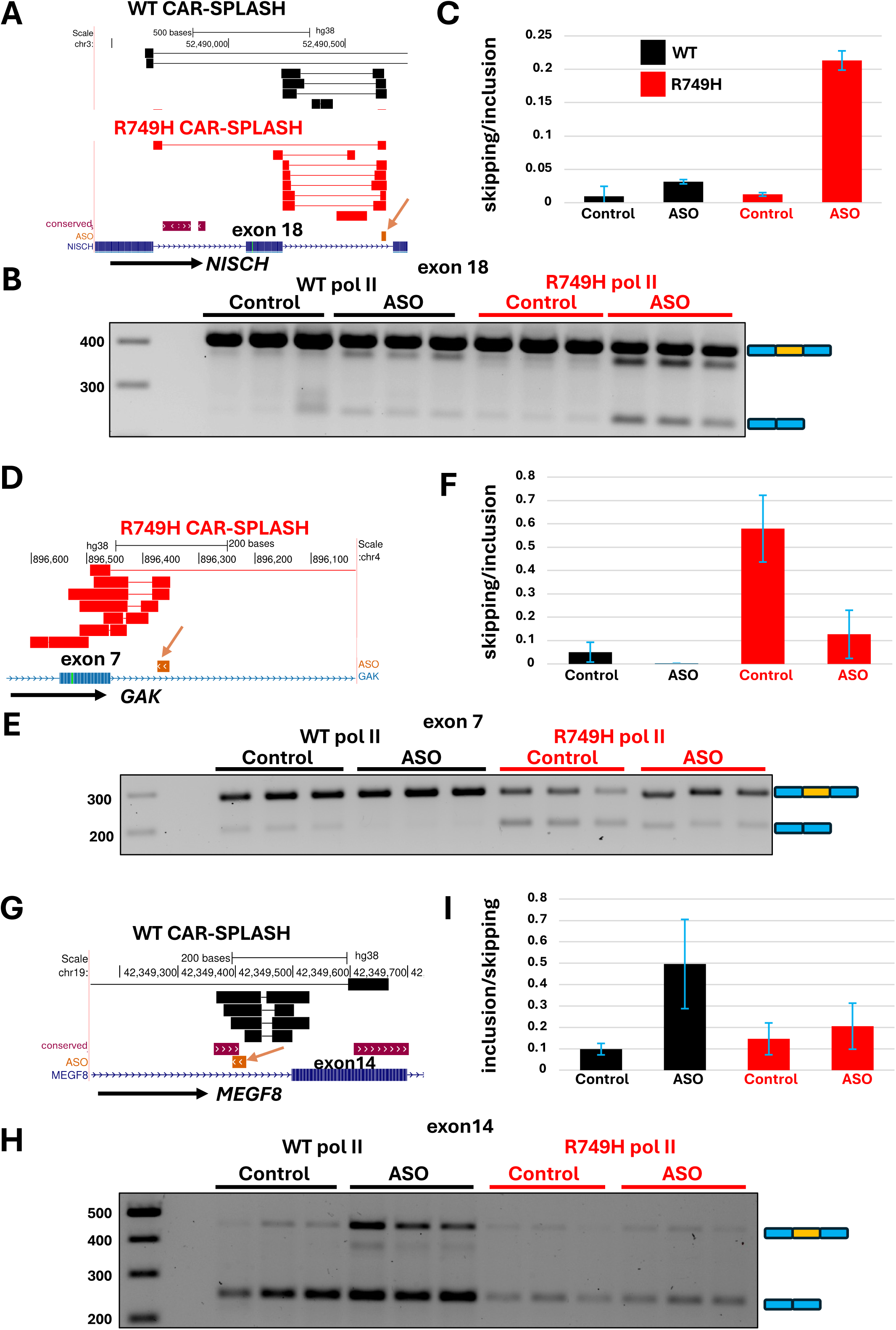
RNA kinetic switches implicated in elongation rate dependent splicing control. **A.** Genome browser view of CAR-SPLASH hybrid reads in cells expressing WT (black) and slow mutant pol II (red). The ASO (orange arrow, see Supp. Fig. 2B) and conserved RNA structures are indicated. **B, C.** RT-PCR analysis of NISCH exon 18 skipping/inclusion in cells expressing the WT or R749H slow pol II, transfected with specific ASO or control oligonucleotide (3 replicates). Skipping of exon 18 was enhanced by the specific ASO, especially under slow transcription elongation conditions. **D.** CAR-SPLASH RNA structure that sequesters the GAK exon 7 5’ splice site was detected specifically in cells expressing the slow mutant pol II (red). The ASO targeting the downstream arm of the structure is indicated (orange arrow, see Supp. Fig. 2C). **E, F.** RT-PCR analysis of exon 7 skipping/inclusion in cells expressing the WT or R749H slow pol II, transfected with a specific or control ASO (3 replicates). Under slow elongation conditions, ASO treatment strongly reduced exon 7 skipping. **G.** CAR-SPLASH RNA structure detected specifically in cells expressing WT pol II (black) that sequesters the MEGF8 exon 14 3’ splice site. The ASO (see Supp. Fig. 2D) and conserved RNA structures are indicated. **H, I.** RT-PCR analysis of exon 14 inclusion/skipping in cells expressing the WT or R749H slow pol II, transfected with a specific ASO or control oligonucleotide (3 replicates). Note the preferentially elevated exon inclusion under WT transcription elongation conditions when the structure that sequesters the 3’ SS is disrupted by the ASO.

We next interrogated two RNA structures predicted to sequester splice sites involved in transcription rate sensitive alternative splicing of GAK and MEGF8 transcripts (Fig. 4D, G). A structure that was only detected in the slow pol II mutant is predicted to sequester the 5’ ss of GAK exon 7 (Figure 4D, Supplemental Figure 2C).which is skipped more frequently in the slow pol II mutant (Figure 4E). To determine whether this nascent pre-mRNA structure contributes to rate sensitive alternative splicing, we designed an ASO to disrupt it by base pairing with the downstream arm of the duplex (Figure 4D, Supplemental Figure 2C). The specific ASO preferentially increased inclusion of GAK exon 7 relative to the control oligo in the slow pol II mutant relative to WT (Figure 4E, F). We conclude that the 5’ ss sequestering structure identified by CAR-SPLASH in slow pol II mutant cells is physiologically relevant to elongation rate sensitive control of exon 7 inclusion. Specifically, our results suggest this structure forms preferentially when transcription is slow and contributes to the increased skipping of exon 7 under these conditions.

We also investigated a structure detected by CAR-SPLASH that sequesters the 3’ ss of exon 14 in the MEGF8 gene and was only detected in WT pol II cells (Figure 4G, Supplemental Figure 2D). ASO disruption of this structure strongly enhanced inclusion of exon 14 in WT pol II cells but had much less effect in cells expressing the slow pol II mutant where the structure was not detected (Figure 4G-I). We conclude that this 3’ ss sequestering structure identified by CAR-SPLASH functions in control of MEGF8 exon 14 inclusion in an elongation rate sensitive manner.

## Discussion

It is well established that pre-mRNA structure can exert a major effect on splicing efficiency and alternative splicing decisions but the number of documented structures that form *in vivo* and function in splicing regulation is quite limited. In the present study, we addressed this shortcoming by adapting the method of <u>S</u>equencing of <u>P</u>soralen Crosslinked, <u>L</u>igated And <u>S</u>elected <u>H</u>ybrids (SPLASH) to <u>C</u>hromatin-<u>A</u>ssociated nascent pre-m<u>R</u>NAs (Figure 1A). CAR-SPLASH employs an improved biotinylated psoralen analogue (60) for crosslinking and enrichment of crosslinked nascent pre-mRNAs, in addition to a library preparation protocol that incorporates UMIs. We identified ∼6500 putative intramolecular RNA structures within pol II transcripts, some of which are predicted on the basis of sequence conservation (Figure 1C, Table S1). Although only a fraction of pre-mRNA structures formed *in vivo* can be recovered with RNA proximity ligation methods, our CAR-SPLASH data serves as a valuable resource for future investigations, including targeted chemical modification approaches (35, 63, 64, 67, 68). Notably, in this study, we applied stringent criteria to define duplexes with potential roles in splicing regulation, focusing on narrow 200-nt windows relative to splice sites, despite the potential involvement of more distal regions (52). This likely contributed to our success in ASO targeting. However, the set of hybrid reads in our data provides a resource for future studies, where exploring structures with a less restrictive approach could yield further insights into splicing regulation. A total of 673 structures identified by CAR-SPLASH fall within 200 bases of intron-exon boundaries and may therefore influence splicing (Figure 1C, Table S1). Furthermore, many of these lie proximal to alternative splicing events that are sensitive to pol II transcription rate (Figure 3B). These structures are candidate mediators of kinetic coupling between transcription and splicing. We interrogated several of them by transfection of ASOs designed to disrupt structure by base pairing to the arm of the duplex that does not overlap with the splice site. Out of 7 ASO’s targeted to 7 different candidate RNA structures predicted to affect alternative splicing, 4 reproducibly modulated alternative splicing in WT and/or slow pol II mutant expressing cells (Figures 2, 4). We conclude that pre-mRNA structures detected by CAR-SPLASH include bona fide targets for splice modifying ASOs.

Among the structures revealed by CAR-SPLASH, we found three examples in the NISCH, GAK, and MEGF8 pre-mRNAs of a novel type of RNA structural element that we term a “kinetic switch”. These elements affect alternative splicing outcomes in a manner that depends on the RNA pol II elongation rate. We first interrogated a structure identified by CAR-SPLASH that loops out most of intron 18 in the NISCH pre-mRNA. Disruption of this structure with an ASO elicited a previously undetectable isoform that skips exon 18, suggesting that this structure facilitates efficient exon 18 inclusion presumably by juxtaposing intron 18 splice sites, a concept that has been established in model genes but only rarely demonstrated naturally (39, 69). We propose that disruption of this structure increases the effective distance between the 5’ and 3’ ss across intron 18 ultimately leading to increased exon 18 skipping. Notably the NISCH intron 18 ASO had a much greater effect on splicing in the pol II slow mutant than in the WT (Figure 4) suggesting that this structure participates in control of a rate-sensitive alternative splicing event. RNA kinetic switches can also promote exon skipping by sequestering 5’ or 3’ ss in a transcription rate sensitive way. In the case of the GAK exon 7 kinetic switch, slower elongation is predicted to shift the composition of the structural ensemble in favor of 5’ss sequestration, and for the MEGF8 kinetic switch, faster elongation is predicted to favor 3’ ss sequestration. One limitation of the current study is that it lacks the depth to assign relative abundance of RNA structures under WT and slow pol II conditions. Future mechanistic studies of these kinetic switches would benefit from targeted chemical probing methodologies which can reveal the dynamic ensembles of RNA structures (35, 63, 65) that might change under different physiological conditions including those that affect transcriptional elongation.

We suggest that the RNA kinetic switches described here could play a widespread role in mediating kinetic coupling between transcription and pre-mRNA splicing outcomes. Previously splice sites and splicing enhancer and repressor elements have been targeted by ASOs to modify splicing, and this approach has led to therapeutic breakthroughs for treatment of SMN and other diseases (70, 71). The work reported here expands the repertoire of potential targets for therapeutic splice modifiers by showing that pre-mRNA structures detected by psoralen crosslinking can also serve as effective targets for splice modifying ASOs.

## Materials and Methods

### CAR-SPLASH

HEK293 Flp-in WT and slow mutant Rpb1 R749H α-Am^r^ cells have been described (17). α-Am^r^ resistant Rpb1 expression was induced with 2 µg/mL doxycycline for 24 hours followed by a 24 hour incubation with 5 µg/mL α-amanitin and 2 µg/mL doxycycline. Two 15 cm plates at ∼70% confluence were used for each replicate and two replicates were performed for WT and R749H slow mutant cells. Cells were washed with PBS and incubated with 0.2 mM AMT-PEG_3_-Biotin (AP3B), 0.01% digitonin, in PBS for 5 minutes at 37°C as described (1) then crosslinked with 365 nm (2.5 mW/cm^2^) for 20 minutes. Nuclei were isolated and washed with 1M urea, 0.3M NaCl for 15 minutes on ice (61) to enrich for chromatin associated RNA. Chromatin pellets were washed 2X with DNase I buffer (10 mM Tris-HCl pH 7.5, 2.5 mM MgCl_2_, and 0.1 mM CaCl_2_) sonicated with Bioruptor pico and digested with Turbo DNase (0.1U/ µL, 37 °C 1 hr) then with proteinase K (∼1 mg/mL, 1 M GuSCN, 37°C 1 h) followed by Total Nucleic Acid (TNA) extraction, precipitation and re-digestion with Turbo DNase as described (72). Nanodrop and Tapestation HS RNA analysis were used to assess yields and RNA quality.

Chromatin associated RNA was digested with ShortCut RNAse III (0.2 U/µL) in 1X RNAse III buffer (50 mM NaCl, 10 mM Tris-HCl pH 7.9, 10 mM MgCl2, 1 mM DTT) for 7.5 minutes then purified by phenol extraction and crosslinked RNAs were purified by binding to MyOne Streptavidin C1 beads and washed 5X with binding buffer (100 mM Tris-HCl pH 7.5, 10 mM EDTA, 1 M NaCl, 0.1% Tween-20) as described (65).

On Bead Proximity Ligation. Samples were washed with 50 mM Tris-HCl pH 7.5,10 mM MgCl_2_. then resuspended in 1X T4 RNA ligase buffer, 1 mM ATP, 0.5 U/µL Murine RNase inhibitor, 2.5 U/µL T4 RNA Ligase 1 (NEB M0204) and incubated at 16°C for 16 h.

Ligated RNA was eluted (95% formamide, 10 mM Tris-HCl pH 7.5, 10 mM EDTA, 65°C, 5 minutes in a Thermomixer) and precipitated. Eluted RNA was supplemented with Acridine Orange (0.25 mM) and Murine RNase inhibitor (2 U/µL) and reverse crosslinked on parafilm with 254 nm UV (on ice, 4.25 mW/cm^2^, 10 minutes) as described (72). Reverse crosslinked RNA was TNA extracted, precipitated and used as input for the QIASeq UDI miRNA UDI library kit (cat. # 331905) which incorporates UMIs according to the manufacturer’s instructions with the exception that the RT enzyme was substituted with SuperScript IV.

Additional methods are described in the supporting information.

## Supporting information

Intramolecular pre-mRNA clusters of hybrid reads detected by CAR-SPLASH.

## Acknowledgements

We thank Hening Lin and Min Yang (Cornell U) for a generous gift of AP3B, Kent Riemondy and David Farrell for informatics and computational support, and Yue Wan, Zhipeng Lu, Omer Ziv and Andrej Lupták for helpful discussions and Usman Hyder, Jay Hesselberth and Matthew Taliaferro (U Colorado) for comments on the manuscript. We also thank Anob Charkrabarti (Crick Institute) for providing access to *Tosca* prior to publication and Ana Fiszbein (Boston U) for help with RNA seq. This work was supported by the UC Denver RNA Bioscience Initiative and NIH grant 6 to D.B.

---

**Supplemental Figure 1.**
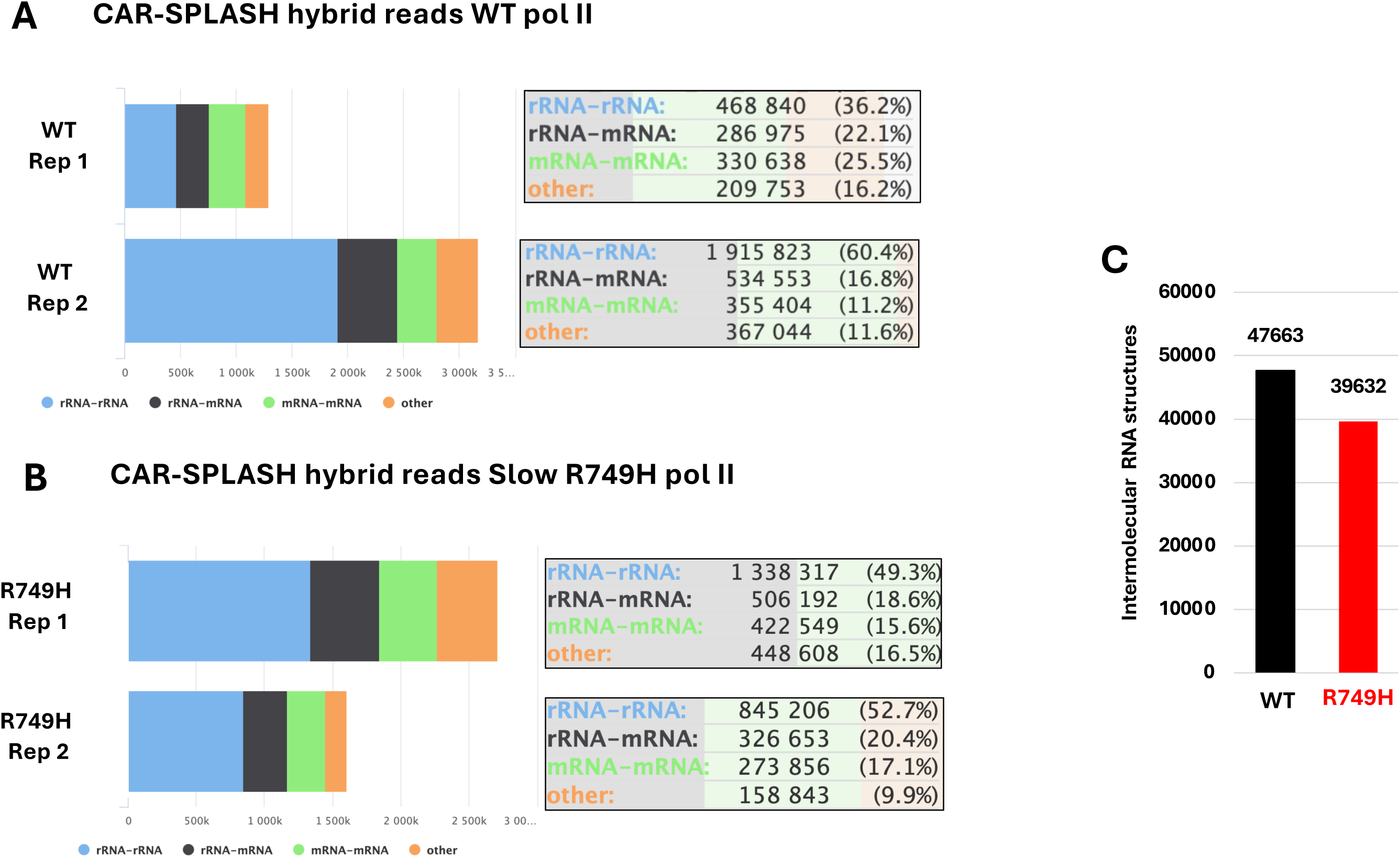
**A, B.** Distribution of CAR-SPLASH hybrid reads between different RNA species as identified by *Tosca* for two replicates each in HEK293 expressing WT and slow mutant (R749H) pol II. **C.** Intermolecular RNA structures detected by CAR-SPLASH in WT and slow pol II mutant cells.

**Supplemental Figure 2.**
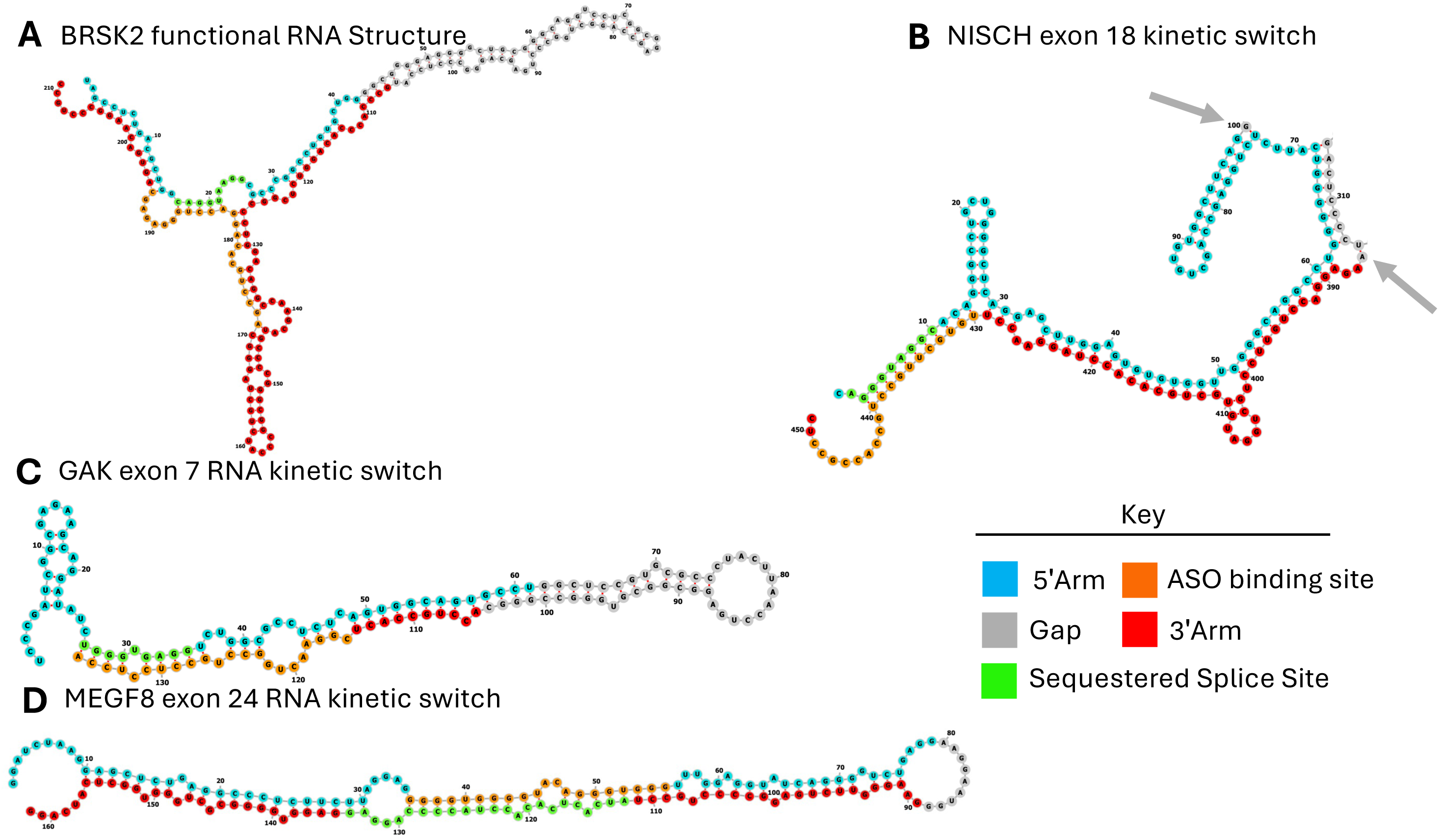
Predicted minimum free energy RNA secondary structures generated using forna (74)(ViennaRNA Web services http://rna.tbi.univie.ac.at/forna/) based on CAR-SPLASH hybrids proximal to alternative splicing events (see Figures 2, 4). The color annotation highlights predicted structural elements: CAR-SPLASH-identified arms, sequestered splice sites, and the ASO-targeted RNA fragment. Grey arrows mark the large loop not shown in the NISCH structure.

